# Immune-Mediated Necrotic Cell Death Initiated by Stressed Cardiomyocytes is a Major Contributor to Cardiomyocyte Loss Following Myocardial Infarction

**DOI:** 10.64898/2026.05.01.722122

**Authors:** Tin Kyaw, Peter Kanellakis, Anne Le, Yi Ee Lye, Pranjal Patel, Kurt Brassington, Nalin Dayawanmsa, Hericka B. Figueiredo Galvao, Grant R. Drummond, Christopher G. Sobey, Alex Bobik, Karlheinz Peter

## Abstract

**Aims:** Percutaneous coronary intervention has improved survival following myocardial infarction, yet strategies to further reduce infarct size are limited. This study investigates the role of cytotoxic γδ-T cells in ischemic cardiomyocyte death and potential therapeutic interventions to reduce infarct size.

**Methods:** Genetic and pharmacological approaches were used to delete γδ-T cells and their specific proteins to assess their involvement in cardiomyocyte death using mouse models of permanent ligation (PL) and ischemia/reperfusion (IR).

**Results:** γδ-T cells accumulated in infarct zones within 6h post-PL, expressing IFN-γ, TNF-α, granzyme B, and perforin. Their deletion reduced infarct size by 73% (PL) and 64% (IR). They induced cardiomyocyte death via apoptosis, gasdermin E-dependent pyroptosis, and MLKL-dependent necroptosis; γδ-T cell depletion reduced apoptosis by 80% and pyroptosis by 38%, with perforin deletion yielding similar effects. Necroptosis, attributed to combined IFN-γ/TNF-α cytotoxicity, decreased by 67%. Cytoplasmic DNA (cDNA) in stressed cardiomyocytes activated the cGAS/STING pathway, inducing expression of chemoattractant MCP-1 and death signal RAE-1. These signals recruited and activated γδ-T cells, which then triggered the death of the stressed cardiomyocytes. STING inhibition suppressed these expressions, reducing γδ-T cell accumulation and infarct size. NKG2D-deficient γδ-T cells prevented activation and reduced infarct size. Administration of an anti-IFNAR antibody at PL onset markedly reduced infarct size.

**Conclusion:** Early activation of cytotoxic γδ-T cells via cardiomyocyte stress signals contributes significantly to immunogenic cardiomyocyte death. Targeting the STING pathway and type I interferon signalling presents a promising therapeutic avenue to mitigate infarct size and improve outcomes.

## INTRODUCTION

Survival following myocardial infarction (MI) has greatly improved in recent decades, largely through the use of percutaneous coronary intervention, but further reductions in infarct size to prevent adverse ventricular remodelling and its consequences remain limited^1^. Consequently, the global burden of ischaemic heart failure is still increasing and is now similar to the combined incidence of breast, prostate and bowel cancers^2^, with an equally similar five-year survival^3^. Current knowledge about the mechanisms responsible for cardiomyocyte death during MI is limited, as most investigators focus on investigating reparative mechanisms rather than initiating mechanisms to minimise adverse ventricular remodelling and heart failure^4^.

The immune system is rapidly activated following MI, exerting both beneficial and detrimental effects on the heart. Neutrophil and monocyte infiltration into the left ventricle (LV) occurs early following MI, with effects varying depending on the timing of infiltration. Neutrophils are initially damaging^5^, but later facilitate healing by stimulating macrophage polarisation to a reparative phenotype^6^. Early infiltrating Ly6C^hi^ monocytes initially contribute to inflammation. Similarly, lymphocytes are also attracted to infarcted myocardium and exert harmful or protective effects^7^. B cells augment ischemic injury by producing CCL7, which enhances Ly6C^hi^ monocyte mobilisation/recruitment to the infarcted heart^8^. CD4+ T cells are also activated following MI^9^, mostly regulatory CD4+ Foxp3+ T cells that beneficially influence healing^10^. In addition, γδ-T cells also accumulate in the infarcted heart and increase infarct size, but mechanisms have not been elucidated^11^. It is not known how these cells are attracted to infarcted myocardium, whether their activation depends on specific cell-cell interactions with ischemia-stressed cardiomyocytes or other downstream mechanisms that increase infarct size. γδ-T cells are mostly activated via innate mechanisms and secrete inflammatory cytokines including IFN-γ, TNF-α and IL-17 as well as cytotoxins such as perforin and granzymes that promote cell death^12, 13^.

Here, we demonstrate that ischaemic cardiomyocytes are not passive bystanders but active instigators of γδ-T cell–mediated immunogenic death during acute MI. Building on our earlier observation that MI triggers rapid mitochondrial DNA (mtDNA) release in cardiomyocytes^14^, we now demonstrate that this surge of mtDNA markedly raises cytosolic DNA (cDNA) levels, which are detected by the cGAS–STING sensor. STING activation initiates IRF3/NF-κB signalling, driving expression of the chemokine MCP-1 and the early responder RAE-1, a ligand for activating receptor NKG2D, on stressed cardiomyocytes. MCP-1 recruits γδ-T cells to the infarct zone, where NKG2D–RAE-1 engagement activates them. Activated γδ-T cells secrete IFN-γ and TNF-α and release perforin/granzyme□B, triggering multiple death programmes, such as GSDME-dependent pyroptosis, MLKL-dependent necroptosis, and apoptosis, in RAE-1-expressing cardiomyocytes, thereby enlarging the infarct size.

This study demonstrates that cGAS-STING activation in stressed cardiomyocytes triggers the recruitment and activation of γδ-T cells, leading to immunogenic cardiomyocyte death. Furthermore, our findings show that inhibiting STING-mediated immune responses in stressed cardiomyocytes with a STING antagonist H-151 or blocking γδ-T cell activity using an anti-IFNAR antibody, significantly reduces infarct size and improves cardiac function. These results suggest that targeting this pathway may represent a novel therapeutic strategy to complement existing revascularisation approaches in MI.

## METHODS

### Animals and Ethics

All mouse strains used were on a C57BL/6 background. NKG2D-deficient (NKG2D^-/-^) mice were obtained from the□Jackson Laboratory (Bar Harbor, ME, USA); perforin-deficient (Pfp^-/-^) and TCRδ–deficient (Tcrd^-/-^) mice were provided by Mark□Smyth (Peter□MacCallum Cancer Centre, Melbourne, Australia). Wildtype (WT) C57BL/6 mice were purchased from the Walter and Eliza Hall Institute (Melbourne, Australia). Ethical approval for all animal procedures was obtained from the Animal Ethics Committee at the Alfred Research Alliance in Melbourne, Australia, and all experiments were conducted at the Precinct Animal Centre located at the Baker Heart and Diabetes Institute.

### Generation of chimeric□mice with γδ-T cell specific deficiency of target proteins

Mixed bone-marrow (BM) chimeras in which γδ-T cells selectively lacked Pfp or NKG2D were generated as described previously^15, 16^. Briefly, 6-week-old male WT recipients received lethal γ-irradiation (11□Gy, split dose) and were reconstituted with 5 × 10^6^ mixed BM cells (80□%□Tcrd^-/-^ plus 20□%□Prp^-/-^ or NKG2D^-/-^). Control chimeras received 80□%□Tcrd^-/-^ and 20□% WT BM cells. Four□weeks later, successful reconstitution and selective Pfp or NKG2D deficiency in γδ-T cells were confirmed by flow cytometry.

### PL or IR surgical procedures

Mice underwent permanent ligation (PL) or ischemia-reperfusion (IR) of the left anterior descending (LAD) artery as described previously^14, 17^. Under ketamine/xylazine/atropine anesthesia (100/10/0.05□mg/kg, i.p.), animals were intubated, ventilated, and a thoracotomy was performed to access the heart. A 7-0 silk suture was placed below the left atrial appendage to occlude the LAD. For IR, the ligature was released after 60□min; for PL, it remained tied. Sham mice underwent the same procedure without LAD occlusion. In selected experiments, mice received anti-TCR□γδ mAb (UC7-13D5, BioXCell; 10□mg/kg, i.v.) 48□h before surgery and fortnightly thereafter; depletion was confirmed by flow cytometry. In therapeutic experiments, animals received either H-151 (0.21□mg/day, i.p. for 4 weeks)^15^ and IFNAR1 mAb (MAR1-5A3, 500□µg at 8 and 48□h post-surgery)^18^ with appropriate control treatment.

### Cardiac function assessment

Cardiac haemodynamics were assessed with a 1.4-F high-fidelity pressure catheter (SPR-839; Millar Instruments, Houston,□TX, USA). Mice were anesthetized by intraperitoneal injection of ketamine (80 mg/kg), xylazine (20 mg/kg), and atropine (0.06 mg/kg), placed supine, intubated, and mechanically ventilated. The catheter was introduced through the right common carotid artery into the ascending aorta and advanced across the aortic valve into the left ventricle. Left-ventricular systolic pressure (LVSP), and the maximal and minimal first derivatives of pressure (dP/dt_max_and dP/dt_min_) were recorded and analysed with ADInstruments software (ADInstruments, Bella Vista, NSW, Australia)^15, 19^. Cardiac function was also assessed by echocardiography using the Vevo 2100 system (FUJIFILM VisualSonics) with an MS-550D transducer (22–55 MHz) in some experiment. Mice were anesthetized with 1–1.5% isoflurane and maintained at ∼450–550 bpm. M-mode images were acquired from the parasternal short-axis view at the level of the papillary muscles. Left ventricular internal diameters at end-diastole (LVEDD) and end-systole (LVESD) were measured using Vevo LAB software. Fractional shortening (FS) and ejection fraction (EF) were calculated using the following formulas^20^:

FS (%) = (LVEDD-LVESD)/LVEDD x100

EF (%) = [(LVEDD)3 – (LVESD)3]/(LVEDD)3 x100

### End-point cardiac tissue and tissue collection

After completing hemodynamic measurements, hearts were arrested in diastole with 15% potassium chloride, and the LV dissected into three equal transverse slices and frozen in OTC at −70°C^17^. Cryosections 6□µm thick were prepared for histology and immunofluorescence, whereas 30□µm sections were used for ROS quantification using dihydroethidium. Some LV tissues were snap-frozen in liquid nitrogen and stored at□–80□°C for later analyses. Peripheral blood, spleen, and inguinal lymph nodes were harvested for immune-cell profiling by flow cytometry.

### FACS Immune cell assessment

Flow cytometry analysis was performed following established protocols^14–16^. Fluorochrome-labelled antibodies (unless otherwise specified, antibodies used in FACS were obtained from BD Biosciences, San Jose, CA) were used to analyse immune cells in the peripheral blood, spleen, and lymphocytes. The antibodies utilized for immune cell analysis included anti-CD19, anti-CD4, anti-CD8, anti-NK1.1, anti-TCRβ, anti-CD314 (NKG2D), and anti-TCRγδ (eBioscience) antibodies. To assess the effects of type I interferon on γδ-T cell cell activation, 100,000 γδ-T cells isolated from C57BL/6 donor mice were stimulated with 1 µg/ml anti-CD3 antibody (clone 145-2C11; BD Biosciences), 10 ng/ml recombinant interferon-α1 (IFNA1) protein (ACROBiosystems), or a combination of both for 5 hours^21^. Cells were then stained with fluorescently labeled antibodies. Data acquisition was performed using a BD LSRFortessa cytometer, and cell surface marker expression was analyzed with BD FACSDiva software.

### Infarct size assessment

Myocardial infarct size in PL and IR injury was quantified by picrosirius–stained sections as described previously^22^. Briefly, serial 5□µm LV transverse sections were imaged with a Nikon microscope fitted with a computer-interfaced colour camera (Optimus□Bioscan□2; Thomas Optical Measurement Systems, USA). Infarct scar and total LV areas were manually delineated on each digital image and expressed as a percentage.

### Immunological staining

Frozen sections were stained with primary antibodies targeting TCR□γδ and CD69 (Abcam), CD27 (BioLegend), IFN-γ and TNF-α (BD Biosciences), phospho-MLKL, granzyme B, GSDME, and MCP-1 (Abcam), α-actinin (Abcam/Bioss), MLKL (Biorbyt), NF-κB p65 (Santa□Cruz), and RAE-1 and NKG2D (R&D□Systems) as previously described^15, 16^. Nuclei were counterstained with DAPI. Dual-color fluorescence staining was performed sequentially, and images were acquired using an Olympus□BX61 fluorescence microscope with CellSens software. For DAB staining, sections were incubated with rabbit anti-GSDME (Invitrogen) or anti-MLKL (Biorbyt), followed by biotinylated secondary antibody, avidin–HRP, and DAB substrate.

### Western blot analysis

LV proteins (20□µg) extracted from ischaemic and non-ischaemic areas were separated on 10% SDS-PAGE gels, transferred to PVDF membranes (Bio-Rad), and probed for GSDME (Invitrogen), N-GSDME (Abcam), MLKL (Biorbyt), phospho-MLKL (Abcam), or β-actin (Sigma). After visualisation with ECL (Bio-Rad) and imaging using a ChemiDoc□XRS+, densitometry was performed in ImageJ v1.53, normalised to β-actin, and expressed relative to sham controls.

### Molecular biology analysis

Total RNA was extracted from aortic arches using the RNeasy Fibrous Tissue Mini Kit (Qiagen), following the manufacturer’s instructions. RNA concentration was measured, and 10□ng of total RNA was used for OneStep QuantiFast SYBR Green RT-PCR (Qiagen), incorporating appropriate housekeeping genes and controls as previously described^14^. The primers used are as follows:

MCP-1 (S) - 5′-CTC AGC CAG ATG CAG TTA ACG-3′,

MCP-1 (AS) - 5′-GGG TCA ACT TCA CAT TCA AAG G-3′;

RAE-1 (S) - 5′-GGC AGA CAG ACA CTC CAG CTT C -3′,

RAE-1 (AS) - 5′-CCA CGA AGC ACT TCA CTT CAT CT-3′.

### Statistical Analyses

Results are presented as mean +/- SEM. Comparisons between groups were determined by two tailed Student’s *t-*test and three or more groups were compared by one-way ANOVA followed by Turkey-Kramer *post hoc* analysis. P < 0.05 was considered statistically significant.

## RESULTS

### CD27⁺ γδ-T cells drive cardiomyocyte apoptosis and inflammatory necrosis following myocardial injury

To elucidate the mechanisms by which recruited γδ-T cells contribute to myocardial injury, we first assessed their accumulation and molecular characteristics within the ischemic myocardium. γδ-T cells accumulated significantly in the infarcted regions compared to non-ischemic areas and were detectable as early as 6 hours post-PL and continued to increase in numbers over 5 days (**Figure 1A**). The majority of these cells exhibited high activation status (**Figure 1B**), as indicated by CD69 expression^23^. Phenotypic analysis revealed that most infiltrating γδ-T cells were CD27⁺ (**Figure 1C**), a marker distinguishing IFN-γ-producing γδ-T cells from IL-17-producing subsets^24^. Further analysis revealed that γδ-T cells expressed high levels of IFN-γ and TNF-α (**Figure 1D-E**), cytokines known to mediate inflammatory cell death^25, 26^, as well as cytolytic proteins Pfp and GrzB (**Figure 1F-G**). The expression of these molecules by γδ-T cells suggests their capacity to induce both apoptotic and inflammatory necrotic cell death pathways in ischemic cardiomyocytes. TNF-α and IFN-γ can act synergistically to promote inflammatory cell death^25^, while Pfp and GrzB can trigger either apoptosis or inflammatory cell death^27^.

**Figure 1.**
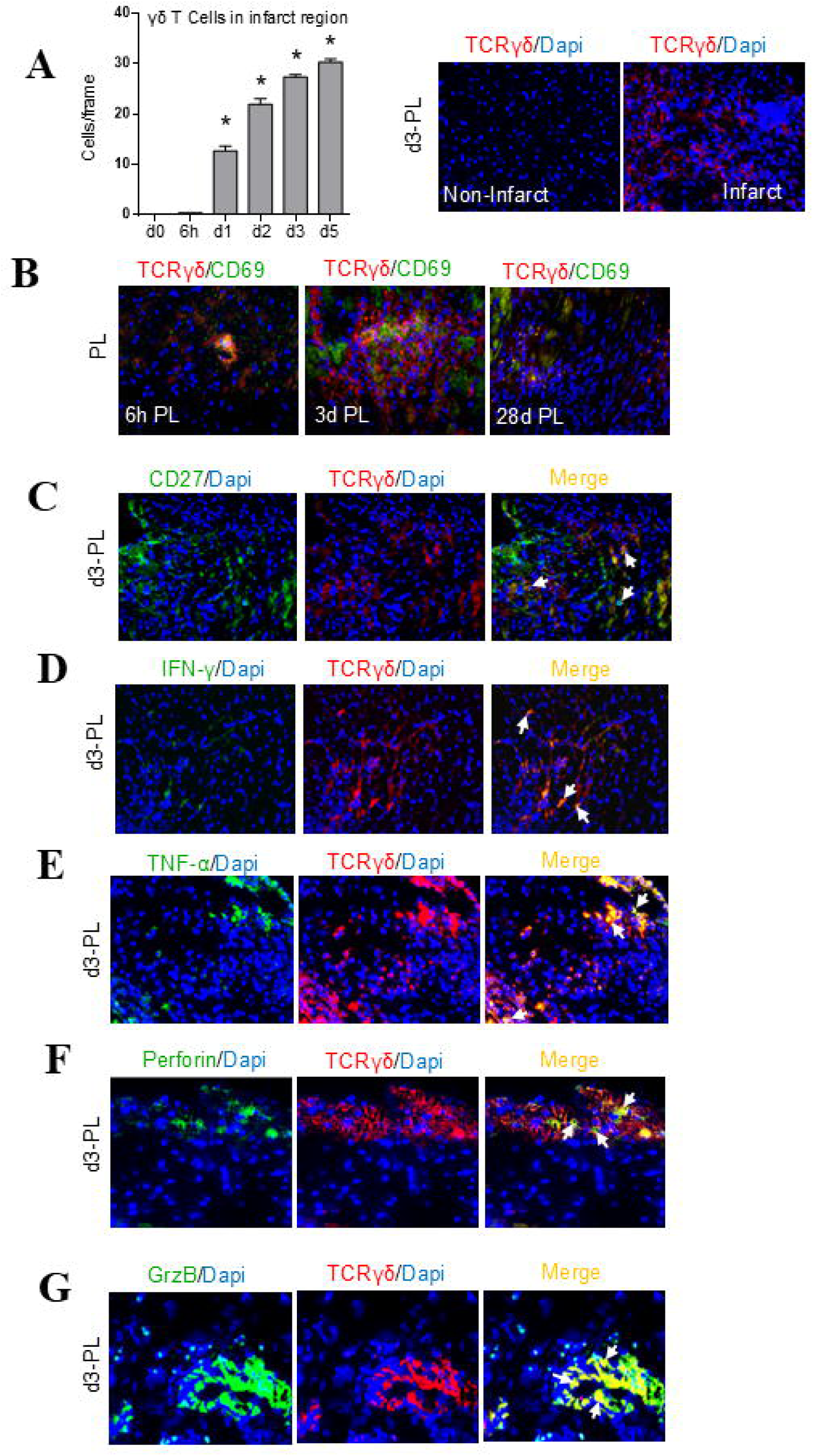
Activated γδ-T cells expressing inflammatory and cytotoxic molecules accumulate in infarcted myocardium. C57Bl/6 mice were subjected to myocardial ischaemia by permanent ligation (PL) of the left anterior descending artery. Ischaemic myocardial tissues were immunofluorescently stained with various antibodies. A significant accumulation of γδ-T cells was observed in the infarcted myocardium 24 hours after PL, and this level remained elevated until day 5 (A). The majority of γδ-T cells expressed the activation marker CD69 (B). Many γδ-T cells also expressed the costimulatory receptor CD27 (C). γδ-T cells in the infarct zone expressed IFN-γ (D), TNF-α (E), Pfp (F), and GrzB (G). Representative photomicrographs are shown from groups (n = 3–7). Data are presented as mean ± S.E.M. p < 0.05 compared to day 0.

### Depletion of γδ-T cells reduces infarct size and improves cardiac function

To assess the functional impact of γδ-T cells on myocardial injury, we employed a depletion strategy using anti-γδ-T cell antibodies, initiated one day prior to PL. This approach resulted in approximately 95% reduction of γδ-T cells in the spleen, confirmed by the loss of surface and intracellular γδ-T cell receptors (**Figure 2A**). Depletion of γδ-T cells resulted in a significant reduction in infarct size by approximately 65% (**Figure 2B**). The treatment markedly improved left ventricular (LV) function, as evidenced by increased LV□+dP/dt and −dP/dt, a reduced LV-to-body weight (BW) ratio, enhanced fractional shortening (FS), and improved ejection fraction (EF) (**Figure 2C**). To determine the relevance of these findings in the context of reperfusion therapy, we evaluated the effects of γδ-T cell depletion in a mouse model of IR injury. Replacement fibrosis, assessed via picrosirius red staining as an indirect measure of cardiomyocyte loss^28^, was reduced by nearly 60% (**Figure 2D**) following γδ-T cell depletion (**Data not shown**). Correspondingly, improvements in LV function were observed, as assessed by increased LV +dP/dt and −dP/dt, as well as a reduced BW ratio (**Figure 2E**). Collectively, these findings indicate that γδ-T cells accumulating in the ischemic myocardium play a pivotal role in mediating cardiomyocyte death through both apoptotic and inflammatory necrotic pathways following MI, and that their depletion provides significant cardioprotection.

**Figure 2.**
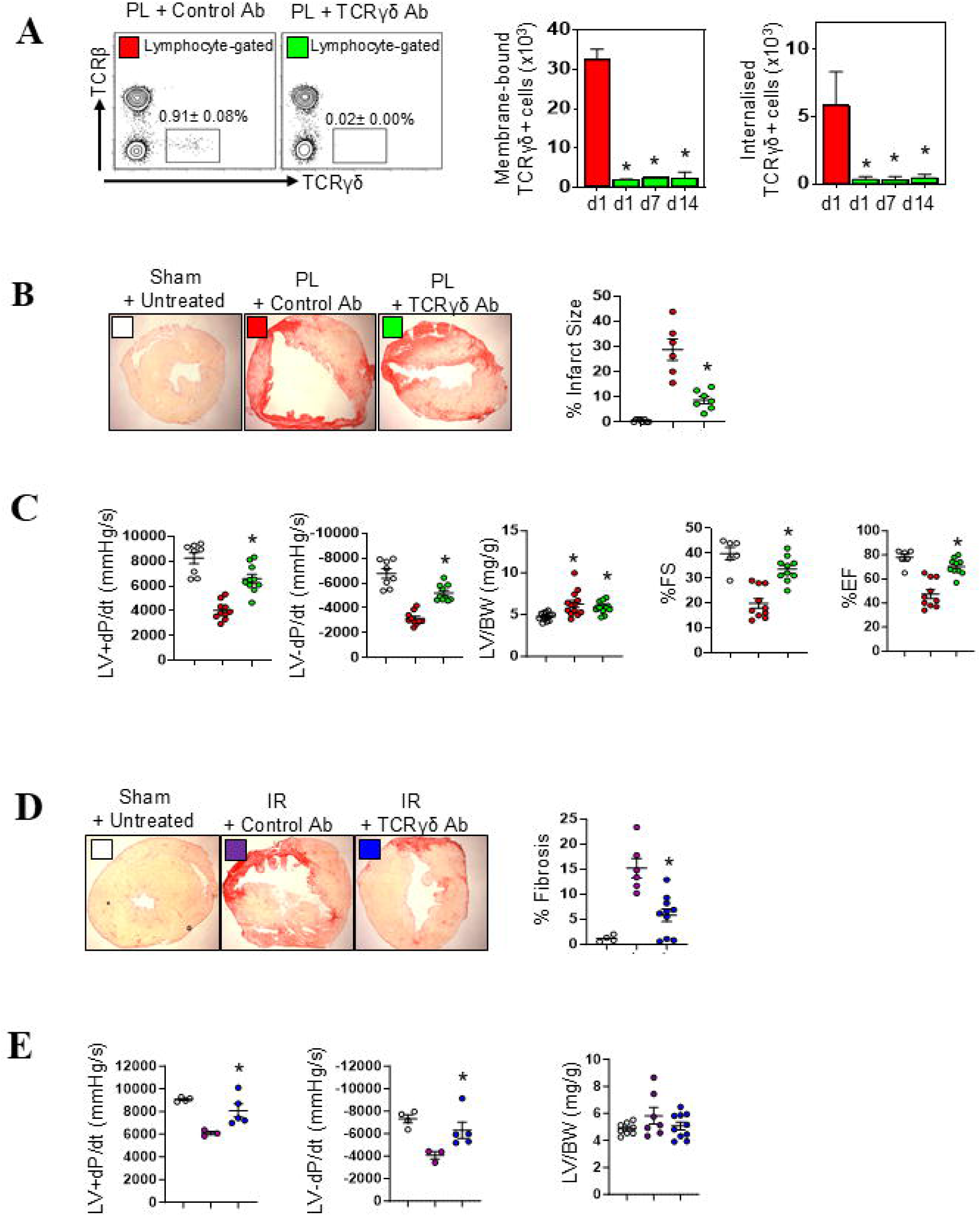
γδ-T cells contribute to infarct size following PL and ischaemia/reperfusion (IR). C57Bl/6 mice pretreated with γδ-T cell–depleting antibodies were subjected to PL or IR injury. Four weeks later, infarct size and cardiac function were assessed. Representative FACS plots obtained two weeks later confirmed near-complete depletion of splenic γδ-T cells following treatment with anti-TCRγδ depleting antibodies (A). In PL-injured mice, depletion of γδ-T cells significantly reduced infarct size (B) and improved left ventricular (LV) function (C). Similarly, in the IR model, γδ-T cell depletion led to markedly smaller infarct size (D) and significantly improved LV function (E). Representative FACS plots and photomicrographs are shown from each group (n = 7–10). Data are presented as mean ± S.E.M. Each dot represents an individual mouse. *p* < 0.05 compared to control antibody–treated group.

### γδ-T cell-derived Pfp induces apoptosis and GSDME-mediated pyroptosis in ischemic myocardium

To elucidate the mechanisms by which γδ-T cells contribute to ischaemic cardiomyocyte death, we investigated their involvement in both apoptotic and pyroptotic pathways. Pfp, expressed by γδ-T cells, is a potent inducer of apoptosis, primarily through the activation of caspase-3^29^. Twenty-four hours after PL, TUNEL assay revealed a significant reduction (∼80%) in cardiomyocyte apoptosis following γδ-T cell depletion (**Figure 3A**). Because lytic forms of cell death, including cytotoxin-induced apoptosis□and□pyroptosis, exacerbate myocardial injury^30^, we assessed the role of γδ-T cells in GSDME-mediated pyroptosis. GSDME, absent in normal left ventricular tissue, was markedly upregulated 24 hours post-PL (**Figure 3B**), likely due to hypoxia-induced epigenetic modifications^31, 32^. Cleaved N-terminal GSDME (N-GSDME), responsible for pore formation and pyroptosis, was abundantly expressed in infarct-zone cardiomyocytes. γδ-T cell depletion significantly attenuated N-GSDME accumulation, indicating their pivotal role in initiating pyroptosis (**Figure 3C**). To confirm the persistence of these effects, we performed Western blot analyses 48 hours post-PL, which demonstrated sustained reductions in N-GSDME levels in the absence of γδ-T cells (**Figure 3D**).

**Figure 3.**
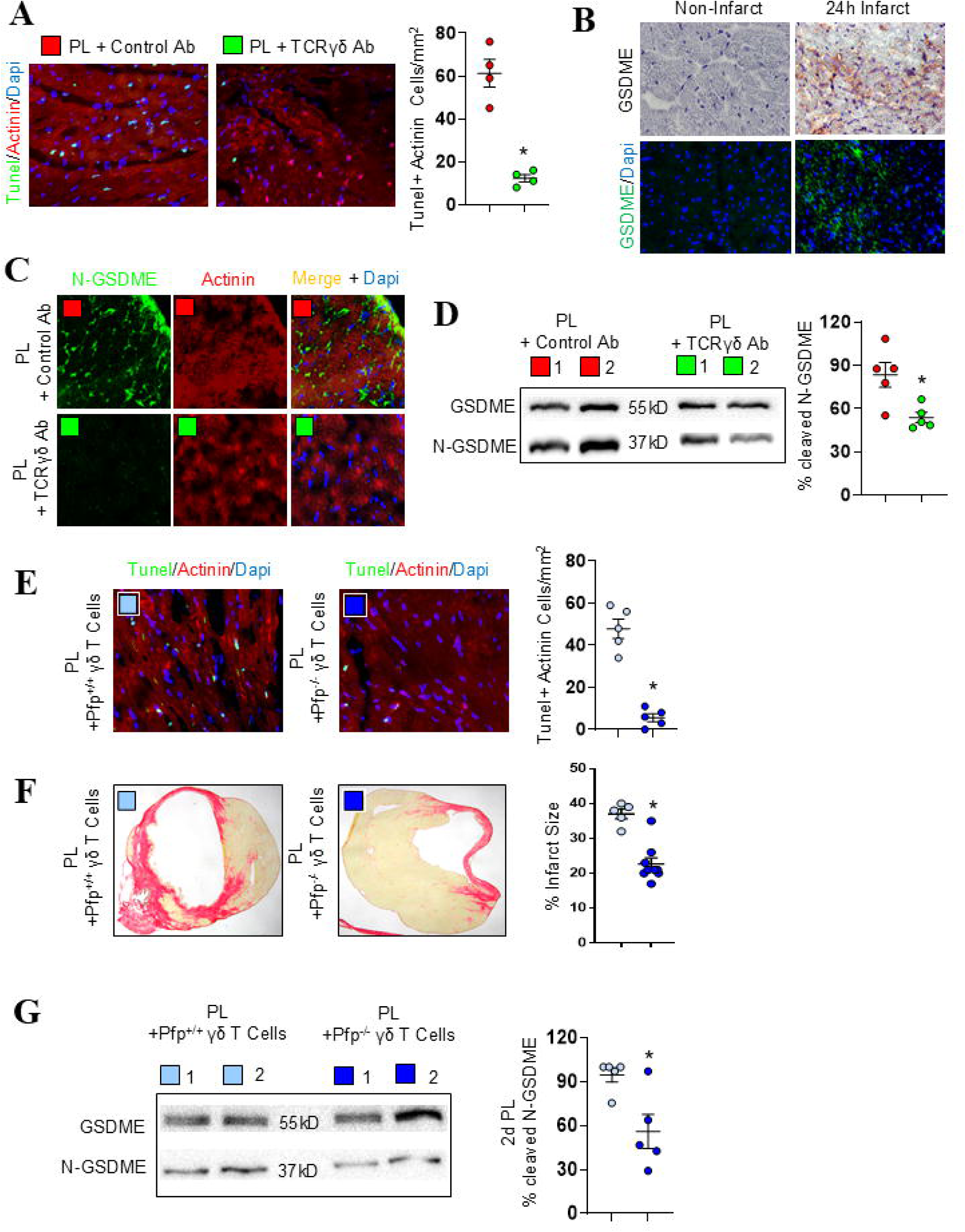
γδ-T cell-derived Pfp promotes apoptosis and GSDME-mediated pyroptosis following PL. In the ischaemic myocardium following PL, γδ-T cell-depleted mice exhibited reduced cardiomyocyte apoptosis (A). GSDME expression was restricted to the left ventricle (LV) within 24 hours post-PL (B). GSDME pore formation, detected using an anti-cleaved N-terminal GSDME antibody, was observed in ischaemic LV cardiomyocytes and found to be highly dependent on the presence of γδ-T cells, as demonstrated by immunofluorescence and Western blot analyses (C, D). Chimeric mice with γδ-T cell-specific Pfp deficiency showed that cardiomyocyte apoptosis post-PL was critically reliant on γδ-T cell-derived Pfp (E). Loss of Pfp in γδ-T cells significantly reduced infarct size (F) and attenuated GSDME cleavage and pore formation (G). Representative photomicrographs and Western blots are shown for each group. Each dot represents an individual mouse. Data are presented as mean ± S.E.M. p < 0.05 compared to the respective control group.

To directly implicate γδ-T cell-derived Pfp in mediating apoptosis and pyroptosis, we employed a mixed bone marrow chimera model to selectively delete Pfp in γδ-T cells, as we described before^15, 16, 33^. Pfp facilitates the delivery of GrzB into target cells^34^. Pfp-deficient γδ-T cells exhibited an 88% reduction in their ability to induce cardiomyocyte apoptosis (**Figure 3E**) and a nearly 40% decrease in infarct size (**Figure 3F**). Correspondingly, N-GSDME levels in the infarct zone were reduced by approximately 40% (**Figure 3G**), corroborating the role of γδ-T cell-derived cytotoxic granules in promoting both apoptotic and pyroptotic cardiomyocyte death. Collectively, these findings underscore the critical contribution of γδ-T cells to myocardial injury through the induction of apoptosis and GSDME-mediated pyroptosis, highlighting potential therapeutic targets to mitigate infarct size and preserve cardiac function.

### γδ-T cells promote MLKL-mediated necroptosis in infarcted cardiomyocytes

Interferons released from highly stressed cells transcriptionally up-regulate mixed-lineage kinase domain-like protein (MLKL), and TNF-α potently activates MLKL signalling^35, 36^. To determine whether IFN-γ- and TNF-α-producing γδ-T cells drive necroptotic cardiomyocyte death after myocardial ischemia, we quantified MLKL expression and activation. MLKL was undetectable in healthy left-ventricular tissue but became markedly up-regulated 24□h after PL (**Figure 4A**), coincident with the rise in IFN-γ and TNF-α from infiltrating γδ-T cells (**Figure 1**). Phosphorylated MLKL (p-MLKL), a marker of activation^37^, accumulated in cardiomyocytes within the infarct zone and declined significantly when γδ-T cells were absent (**Figure 4B&C**). This reduction persisted at 48□h, with >60□% of p-MLKL-positive cells depending on γδ-T cells (**Figure 4D**). MLKL executes necroptosis downstream of receptor-interacting protein kinase-3 (RIPK-3)^38^. Although pyroptosis and necroptosis arise from distinct pathways, there is evidence to suggest that they may be interlinked^39^. Accordingly, co-staining revealed that ∼15–20□% of dying cardiomyocytes co-expressed p-MLKL and GSDME, indicating overlap between the two lytic programmes (**Figure 4E**). As apoptosis was relatively infrequent compared to pyroptosis and necroptosis, we did not investigate panoptosis, a coordinated cell death program dependent on AIM2, pyrin, and ZBP1, which is more characteristic of virally infected macrophages^40, 41^. Together, these data identify γδ-T cells as pivotal in triggering MLKL-mediated necroptosis in ischemic cardiomyocytes, thereby amplifying inflammatory cell death and infarct size expansion.

**Figure 4.**
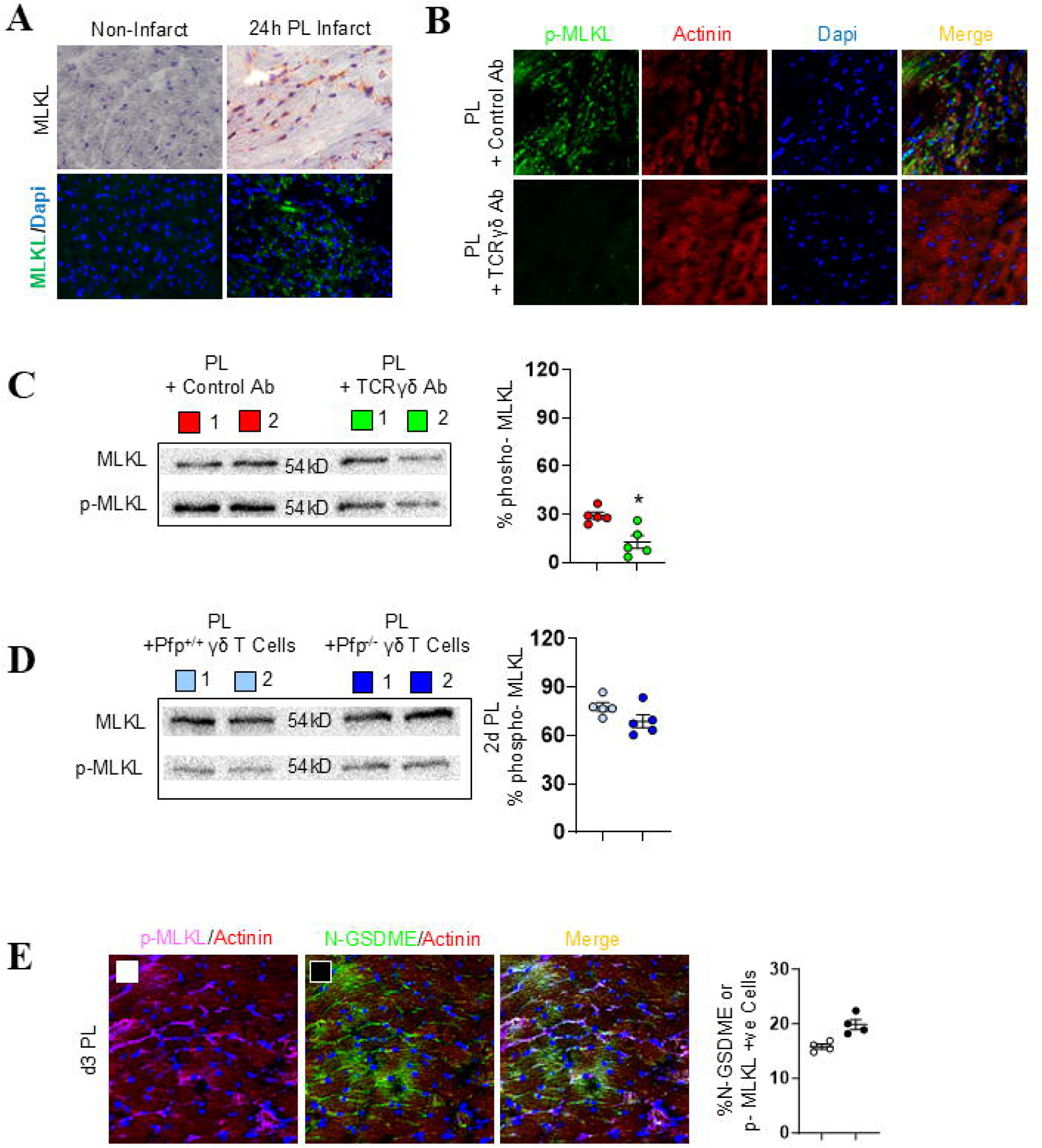
γδ-T cells promote MLKL-mediated necroptosis following PL. MLKL, which is not expressed in healthy left ventricle (LV) tissue, becomes upregulated in the ischaemic myocardium within 24 hours after PL (A). MLKL pore formation in cardiomyocytes, detected using anti-phospho-MLKL antibodies, was highly dependent on the presence of γδ-T cells (B, C). In contrast, γδ-T cell-derived Pfp did not influence MLKL phosphorylation or pore formation (D). Approximately 20% of cardiomyocytes in the infarct zone exhibited features of both pyroptosis and necroptosis, as indicated by co-expression of cleaved N-terminal GSDME and phospho-MLKL (E). Representative photomicrographs and Western blots are shown for each group. Each dot represents an individual mouse. Data are presented as mean ± S.E.M. p < 0.05 compared to the respective control group.

### cGAS–STING activation in infarcted cardiomyocytes drives γδ-T cell recruitment and activation through MCP-1 and RAE-1

Myocardial ischaemia releases mtDNA and nuclear DNA, key triggers of sterile inflammatory cell death^42^. The ensuing surge in cDNA activates the cyclic GMP–AMP synthase–stimulator of interferon genes (cGAS–STING) pathway^15, 43^. We previously showed that cDNA-induced cGAS–STING signalling in cardiac fibrosis up-regulates cardiomyocyte MCP-1 and the stress-induced ligand RAE-1^41^.

Here, cDNA concentrations rose sharply within 2□h of PL onset and continued to increase (**Figure 5A**), consistent with mitochondrial dysfunction^44^. Reactive-oxygen-species (ROS) levels increased in parallel (**Figure 5B**), further indicating mitochondrial impairment^45^. STING activation was confirmed by elevated phospho-STING (p-STING) 24□h post-PL (**Figure 5C**) and by activation of its two downstream branches, phosphorylated IRF3 (p-IRF3) and nuclear NF-κB accumulation (**Figure 5D**). These factors represent two independent pathways activated by STING in stressed cardiomyocytes^15^. NF-κB stimulates chemokine genes such as MCP-1^15, 46^, which is a potent γδ-T -cell chemoattractant^47^, and infarct-zone cardiomyocytes displayed strong MCP-1 expression (**Figure 5E**).In addition, IRF3-dependent RAE-1 was undetectable in healthy hearts but became robustly expressed in infarcted cardiomyocytes (**Figure 5F**), providing the NKG2D ligand necessary for contact-dependent γδ-T -cell activation. Collectively, these findings demonstrate that cDNA-triggered cGAS–STING signalling in ischaemic cardiomyocytes induces MCP-1 and RAE-1, thereby recruiting γδ-T cells and priming them via RAE-1-NKG2D interactions to mount cytotoxic responses that exacerbate infarct injury.

**Figure 5.**
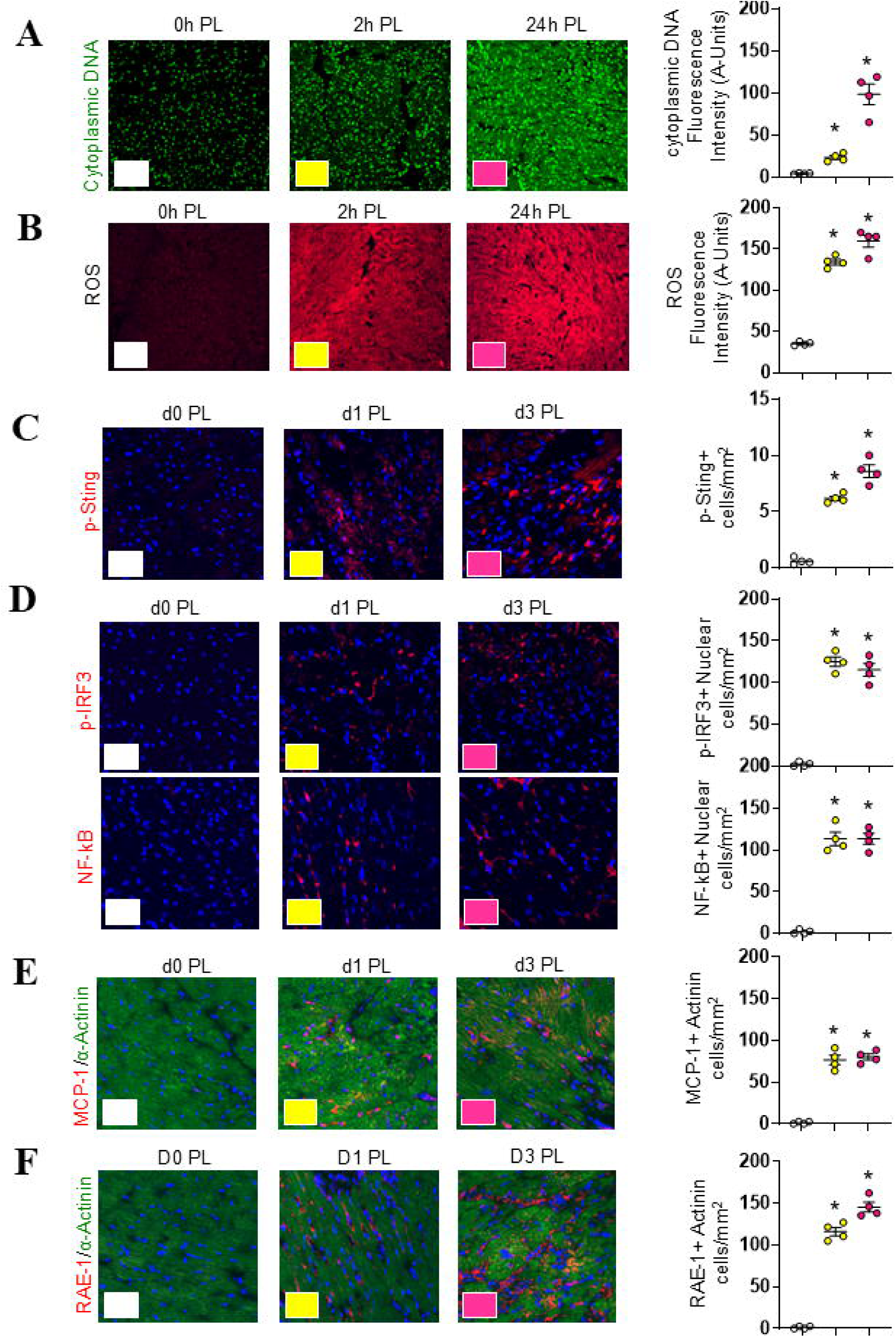
Cell stress signalling upregulates cardiomyocyte expression of MCP-1 and RAE-1 following PL. In the ischaemic myocardium following PL, cDNA levels (detected using Pico488) and reactive oxygen species (ROS) levels (measured with dihydroethidium, DHE) were rapidly increased and remained elevated (A&B). Similarly, STING phosphorylation, detected using an anti-phospho-STING antibody, was elevated in ischaemic cardiomyocytes after PL (C) and was accompanied by increased expression of phosphorylated IRF and nuclear accumulation of NF-κB (D). Within 24 hours following PL, ischaemic cardiomyocytes expressed elevated levels of the chemokine MCP-1 and the NKG2D ligand/stress protein RAE-1, and these levels remained high (E&F). Representative photomicrographs are shown for each group. Each dot represents an individual mouse. Data are presented as mean ± S.E.M. *p* < 0.05 compared to the respective control group.

### IR triggers the cDNA–cGAS–STING Axis, and NKG2D–RAEIZ1 interactions between recruited γδ-T cells and stressed cardiomyocytes regulate infarct size

To verify that the cDNA-cGAS-STING signalling cascade seen after PL is also activated by IR, we subjected mice to 90□min of coronary occlusion followed by reperfusion. Twenty-four hours later, cDNA levels and ROS production were markedly elevated and remained high for ≥3□days (**Figure 6A**).So were p-STING and the stress ligand RAE-1 (**Figure 6B**), consistent with ROS-mediated mitochondrial DNA damage driving cDNA accumulation^48, 49^. Building on our finding that stressed cardiomyocytes up-regulate MCP-1 and RAE-1 after permanent ligation (**Figure 5**), we hypothesise that the same response occurs in the IR model, which better mirrors human MI, thereby triggering death-signal-mediated immune elimination of the stressed cardiomyocytes. To verify that infiltrating γδ-T cells express NKG2D in the I/R myocardium, we colllstained for TCR□γδ and NKG2D. Recruited γδ-T cells showed strong NKG2D expression, and NKG2D frequently co-localized with RAE-1 on stressed cardiomyocytes (**Figure 6C**), indicating the formation of immunological synapses between cytotoxic NKG2D-expressing γδ-T cells and RAE-1–expressing cardiomyocytes, consistent with reports that cytotoxic lymphocytes kill target cells by releasing cytotoxins across the synapse^34^.

**Figure 6.**
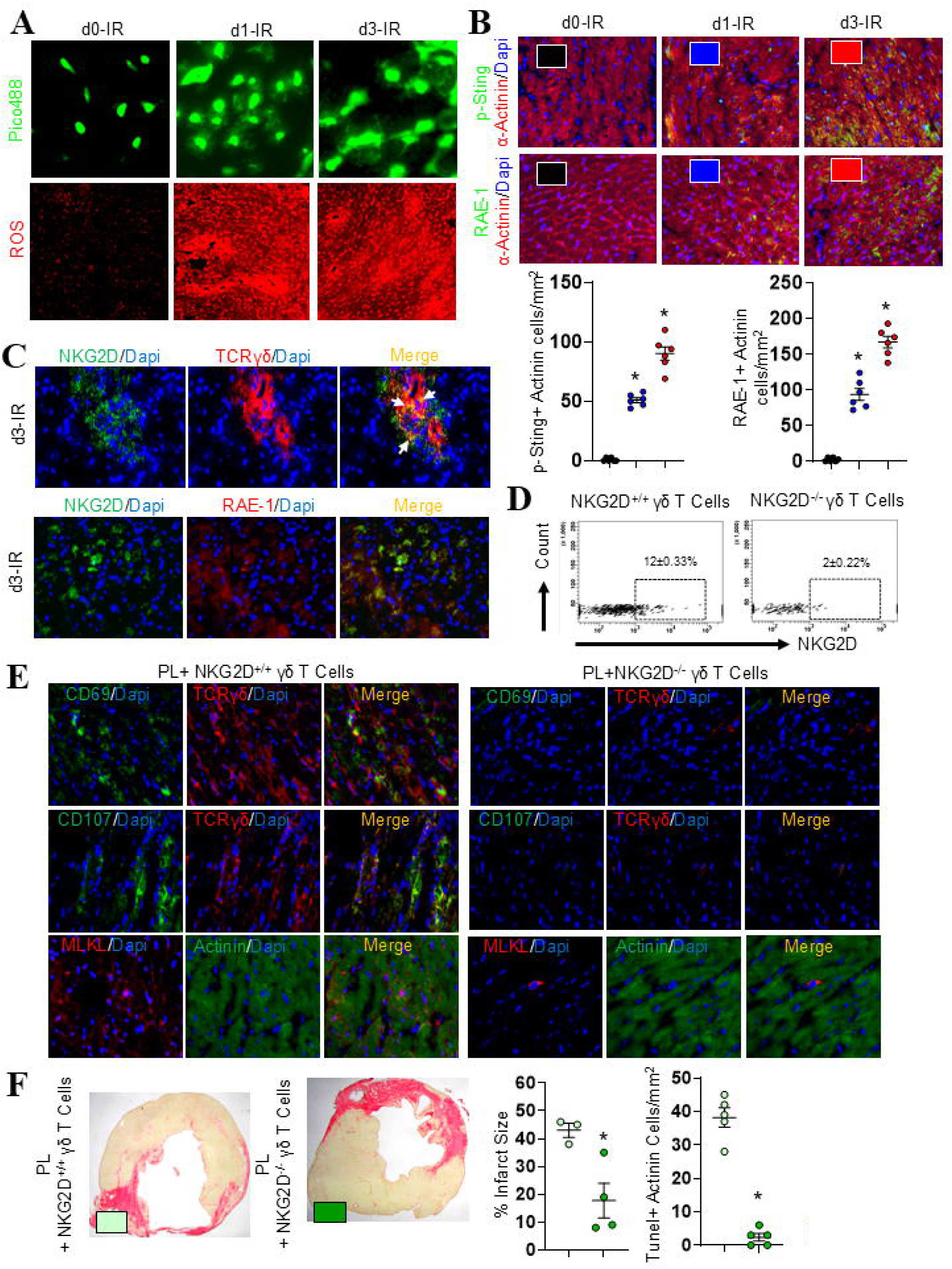
γδ-T cell-specific NKG2D is critical for determining myocardial infarct size. Similar to PL, ischaemia-reperfusion (IR) injury induced large increases in cDNA and ROS levels (A), which were essential for activating STING signaling and upregulating RAE-1 expression within 24 hours, and remained elevated following infarct injury (B). γδ-T cells in the left ventricle (LV) following PL expressed NKG2D receptors, which colocalized with RAE-1 in the infarcted LV (C). Chimeric mice with NKG2D deficiency specifically in γδ-T cells showed complete loss of NKG2D expression (D), which led to impaired γδ-T cell activation (as indicated by reduced CD69 expression), impaired degranulation (CD107a), and attenuated MLKL-mediated necroptosis (E). Deletion of γδ-T cell-specific NKG2D also significantly reduced infarct size and cardiomyocyte apoptosis following PL (F). Representative photomicrographs and FACS dot plots are shown for each group. Each dot represents an individual mouse. Data are presented as mean ± S.E.M. *p* < 0.05 compared to the respective control group.

We next assessed the functional importance of this receptor–ligand interaction and using a mixedlllbonelllmarrow chimera strategy, we deleted NKG2D selectively from γδ-T cells as described before^15, 16, 33^. Flow cytometry confirmed efficient receptor loss (**Figure 6D**). NKG2D deletion abolished γδ-T cell activation after MI, as indicated by the absence of CD69 (**Figure 6E**), and reduced γδ-T cell accumulation by >95□% (**data not shown**). Degranulation capacity, measured by surface exposure of lysosomelllassociated membrane proteinlll1 (CD107a) and IFN–γ–dependent MLKL expression were likewise markedly diminished (**Figure 6E**). Consequently, LV infarct size and cardiomyocyte apoptosis decreased by 60□% and 95% respectively (**Figure 6F**). These findings further demonstrate that NKG2D–RAElll1 engagement between γδ-T cells and stressed cardiomyocytes is a pivotal trigger of immunogenic cardiomyocyte death and a key determinant of infarct size expansion.

### STING inhibition limits infarct size and preserves cardiac function

To assess the functional impact of this pathway on infarct size, we administered the small-molecule STING inhibitor H-151^15^ immediately before PL or before the onset of reperfusion. Twenty-eight days later, left ventricular infarct size was reduced by 88% in the PL model and 63% in the IR model, with both groups of H-151–treated mice showing improved left ventricular function, as indicated by increased ejection fraction and fractional shortening (**Figure 7A&B**). At 3□days post-IR, cardiomyocyte MCP-1 and RAE-1 expression were suppressed by 78□% and 84□%, respectively (**Figure 7C&D**), These results confirm that IR activates the same cDNA–cGAS–STING cascade as PL and demonstrate that pharmacological STING inhibition effectively blunts downstream chemokine/ligand induction, γδ-T cell recruitment, and infarct size expansion. Our data demonstrate that robust STING activation drives MCP-1 and RAE-1 expression in cardiomyocytes during MI (**Figure 4**). As upregulation of MCP-1 in the ischaemic myocardium may be critically required for RAE-1– and NKG2D-mediated elimination of stressed cardiomyocytes, we examined their expression over time following myocardial injury. RT-PCR analysis revealed a rapid and robust upregulation of MCP-1 mRNA in the infarcted myocardium, peaking at day 1 post-MI (>90-fold vs. non-infarct areas), remaining elevated at day 3, and returning to baseline by day 7 (**Figure 7E**). RAE-1 mRNA also rose sharply at day 1 (∼15-fold), stayed elevated at day 3 (∼12-fold) and day 7 (∼8-fold), and returned to baseline by day 14 (**Figure 7F**). No significant changes were observed in non-infarct regions, except at day 1, where both MCP-1 and RAE-1 were modestly elevated, likely reflecting a systemic immune response to MI. Moreover, STING activation generates abundant type□I interferons (IFN-Is), a canonical output of this pathway^50, 51^. Consistent with previous reports demonstrating abundant expression of IFN-Is at sites of myocardial injury, we confirmed their upregulation in ischaemic myocardium compared to non-ischaemic heart tissue (**Figure 7G**). Collectively, MCP-1, RAE-1, and type I interferons (IFN-Is), produced via activation of the cGAS–STING pathway, render stressed cardiomyocytes to act as active instigators of γδ-T cell–mediated immunogenic cell death during acute MI. These findings highlight IFNAR blockade as a potential therapeutic strategy to interrupt this pathogenic cascade and limit myocardial injury.

**Figure 7.**
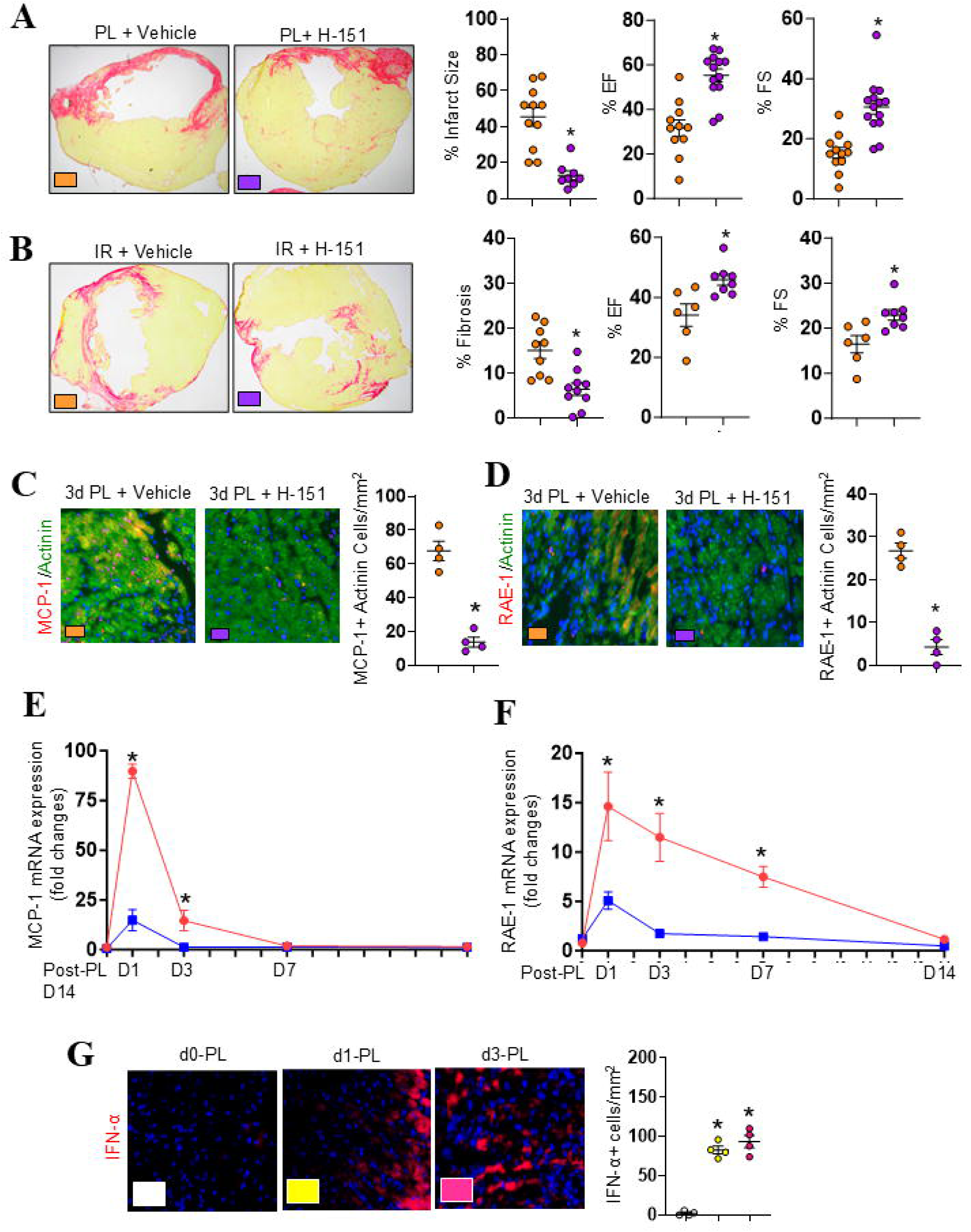
Inhibition of STING signaling reduces infarct size following ischaemic injury. Treatment of mice with the STING inhibitor H-151 significantly reduced infarct size and improved cardiac functions following both PL (A) and IR (B). This treatment was also associated with marked reductions in cardiomyocyte expression of MCP-1 (C) and RAE-1 (D). Using RT-PCR, infarcted myocardium (red line) showed a time-dependent increase in MCP-1, and RAE-1 mRNA expression compared to the non-infarct area (blue line) (E&F). Immunofluorescence analysis revealed upregulation of type I interferon in the infarcted area following PL-induced ischemic injury (G). Representative photomicrographs are shown for each group. Each dot represents an individual mouse. Data are presented as mean ± S.E.M. *p* < 0.05 compared to the respective control group.

### Therapeutic blockade of IFN-I signalling in γδ-T cells limits infarct size and improves cardiac function

IFN-I signalling serves as a pivotal extrinsic cue for γδ-T cells, enhancing their IFN-γ production and cytotoxic responses during infections and tumors^52, 53^. However, in certain viral infections, IFN-I signalling can suppress IL-17 production by γδ-T cells^54^, thereby dampening their pro-inflammatory responses. To delineate the specific effects of IFN-I signaling on γδ T-cell activation, recombinant IFNA1 was used in an in vitro stimulation assay, either alone or in combination with anti-CD3 antibody^21^, as γδ T-cell responses are known to be potentiated by TCR activation, a dominant feature of myocardial infarction^55^. Flow cytometry analysis demonstrated that γδ-T cells stimulated with both IFNA1 and anti-CD3 antibody exhibited a marked increase in CD69 expression compared with untreated or singly treated cells (**Figure 8A**), underscoring the critical role of IFN-I signaling in enhancing γδ T-cell activation.. These findings suggest that γδ-T cells recruited by MCP-1 and activated by IFN-Is at sites of myocardial injury are equipped to eliminate RAE-1–expressing stressed cardiomyocytes via a NKG2D-dependent cytotoxic mechanism. To assess whether interrupting this pathological loop confers cardioprotection, we treated mice with a monoclonal antibody targeting IFNAR1 in a clinically relevant IR model. Four weeks post-treatment, IFNAR1 blockade significantly reduced infarct size (**Figure 8B**) and improved left ventricular function (**Figure 8C-D**). Notably, anti-IFNAR1 mAb treatment also decreased the accumulation of CD69⁺ γδ-T cells in infarcted myocardium (**Figure 8E**), supporting a therapeutic mechanism mediated via γδ-T cells rather than direct effects on stressed cardiomyocytes.

**Figure 8.**
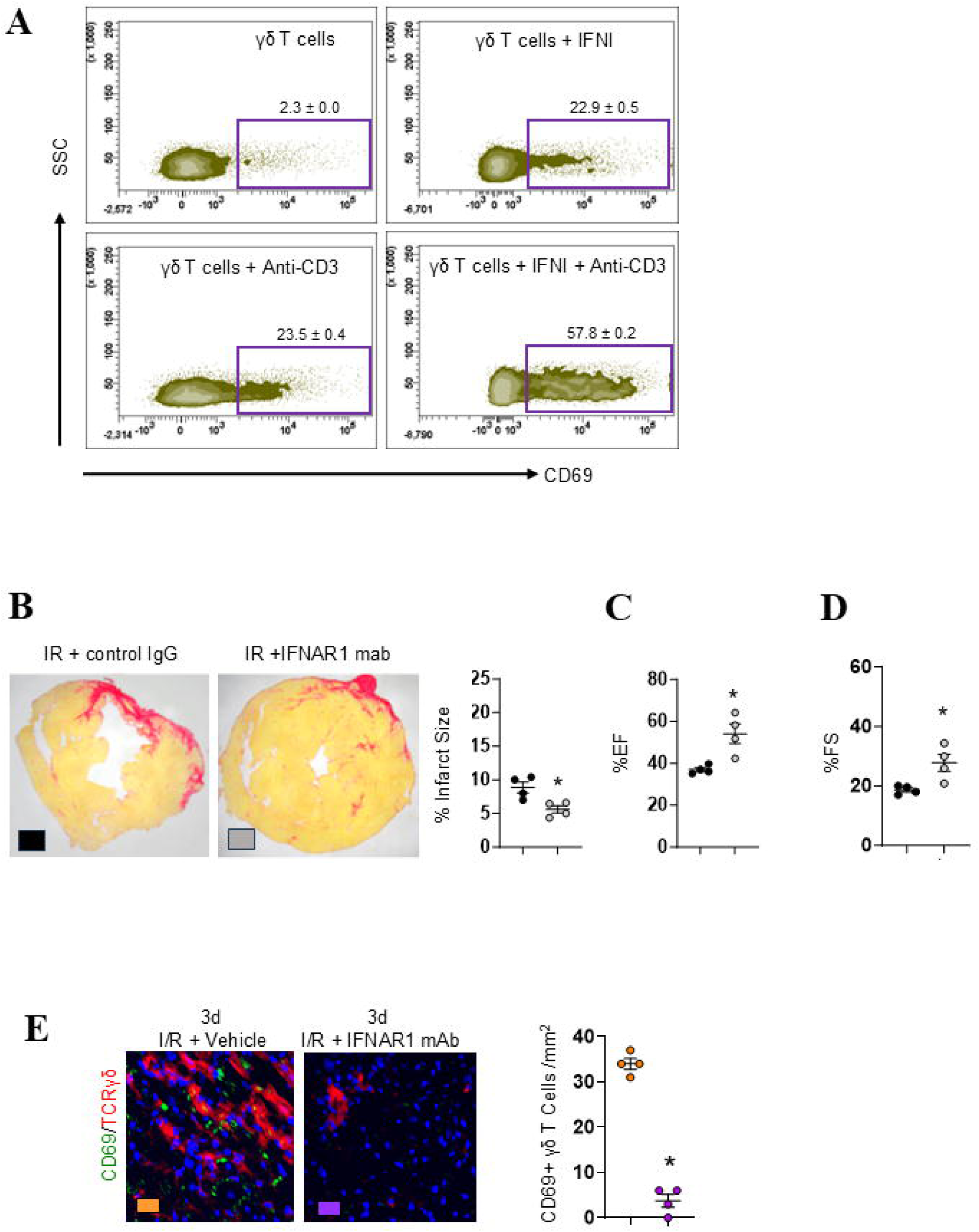
Type I IFN is critical for γδ T-cell activation, and IFN-I blockade reduces infarct size. γδ-T cells were stimulated in vitro with anti-CD3 antibody, IFNA1, or both. Combined stimulation markedly increased CD69 surface expression compared with untreated or singly treated cells, indicating that type I interferon promotes γδ T-cell activation (A). Treatment with an IFNAR1-blocking antibody significantly reduced infarct size (B), ejection fraction (C) and fractional shortening (C), reduced CD69+ γδ-T cells in infarcted areas (E). Representative FACS plots and photomicrographs are shown from each group. Each dot represents an individual mouse. Data are presented as mean ± S.E.M. p < 0.05 compared to the respective control group.

## DISCUSSION

Our results identify an innate immune surveillance system responsible for monitoring cell health^56, 57^ as a key driver of immunogenic cardiomyocyte death in both PL and IR injury. Shortly after the onset of ventricular ischemia, stressed cardiomyocytes accumulate cDNA, activating the cGAS–STING pathway. STING signalling up-regulates secretory chemokines, such as MCP-1, and membrane-bound stress signals, such as RAE-1. Together they recruit and prime cytotoxic γδ-T cells, early responders to infections, injuries and inflammation, to eliminate stressed cardiomyocytes through both inflammatory and non-inflammatory mechanisms that critically depend on NKG2D–RAE-1 engagement within immune synapses. Finally, we demonstrate that transient blockade of the STING pathway using the STING antagonist H-151, or inhibition of type□I interferon signalling, a key downstream effector of STING, via an anti-IFNAR antibody, markedly reduces infarct size. This immunomodulatory strategy represents a promising adjunct to current revascularisation therapies for mitigating the adverse consequences of myocardial injury.

Ischaemic cardiomyocytes are traditionally thought to die primarily from metabolic and ionic derangements^58^; yet the upstream molecular triggers remain ill⍰defined. Prompted by evidence that cDNA serves as a danger signal, we investigated its contribution to cardiomyocyte death in both PL and IR models. We previously reported that MI provokes the rapid release of mtDNA^14^, a strongly immunogenic molecule capable of eliciting broad inflammatory responses^59^. Here we show that organelle⍰derived DNA appears in the cardiomyocyte cytoplasm as early as 2□h post-MI. We next asked whether this early surge in cDNA acts as a “distress signal” analogous to those displayed by virally infected cells^60^, thereby prompting cardiomyocytes to present immune⍰activating surface ligands. Although mtDNA is recognized as ‘foreign’, the cGAS–STING sensor also detects mis⍰localised nuclear DNA once it gains cytosolic access^60, 61^. In the present study we quantified total cDNA, including both nuclear and mitochondrial sources, while our earlier human data confirmed mtDNA release after MI^14^. Elevated cDNA closely correlated with activation of the cGAS–STING pathway, evidenced by increased p⍰STING and its downstream effectors p⍰IRF3 and NF⍰κB. These transcription factors drive expression of the chemokine MCP⍰1, the NKG2D ligand RAE⍰1, and type□I interferons, thereby linking cDNA accumulation to the immunogenic elimination of ischaemic cardiomyocytes^62–64^.

Pharmacological STING inhibition markedly attenuated injury in both PL and IR models, an effect accompanied by sharp reductions in cardiomyocyte MCP-1 and RAE-1 expression. Numbers of activated γδ-T cells within the infarct zone likewise decreased, consistent with stressed-cell death being driven by RAE-1–NKG2D interactions. In human MI, the NKG2D ligand MICA is markedly elevated in plasma following myocardial injury^65^, and similar to murine RAE-1, its expression is induced by DNA damage^66^. Whether other NKG2D ligands are also elevated after MI remains to be determined; in mice, ligands such as MULT1 are more closely associated with endoplasmic-reticulum stress^67^ whereas several other human stress proteins are typically up-regulated in oncogenically transformed cells^68^.

To delineate how γδ-T cells provoke immunogenic cardiomyocyte death, we profiled the cytotoxic mediators they express in the infarct zone. Infiltrating γδ-T cells displayed a canonical cytolytic signature, such as Pfp, GrzB, IFN-γ and TNF-α, suggesting these effectors act during the cell-to-cell contacts we observed between γδ-T cells and stressed cardiomyocytes. Ablating γδ-T cells markedly reduced cardiomyocyte apoptosis and, unexpectedly, attenuated GSDME-dependent pyroptosis, even though healthy myocardium lacks GSDME. GSDME first appeared 24□h after MI onset, implicating ischaemic injury in its induction. Indeed, GSDME transcription can be activated by DNA demethylation^69^, a process driven by hypoxia-responsive, oxygen-sensing histone demethylases^31, 32^. Once expressed, GSDME is cleaved by caspase-3 downstream of GrzB^70^; the liberated N-terminal fragment forms membrane pores that culminate in cell lysis. Deleting Pfp in γδ-T cells, which prevents GrzB entry into target cells^71^., mirrored γδ-cell ablation, halving infarct size. Although GSDMD has been recognised as a key pyroptotic effector after MI^72^, our data reveal that it is not acting alone: as in kidney injury^73^, a synergistic interplay between GSDMD and GSDME appears to drive pyroptotic death in the infarcted heart.

Our findings also indicate that γδ-T cell–derived cytokines also drive MLKL-dependent necroptosis in both the PL and IR models; necroptosis requires the combined action of IFN-γ and TNF-α^25^. MLKL is absent from non-ischemic cardiomyocytes, but its expression in infarct-zone cells is induced by interferons^36^. Type□I IFNs are released autocrinally by cGAS-STING-activated cardiomyocytes^34^ whereas type□II IFN (IFN-γ) is secreted by infiltrating γδ-T cells (**Figure 1**). Once MLKL is expressed, TNF-α and IFN-γ promote its phosphorylation and execution of necroptosis^25^. The same cytokine pair also enhances cleavage of GSDMD, GSDME and caspase-3, a molecular signature of panoptosis, a death programme that integrates necroptosis, pyroptosis and apoptosis^25^. Whether this type of death occurs during PL and IR remains to be determined. Immunofluorescent analyses show many infarct-zone cardiomyocytes co-express p-MLKL and cleaved GSDME, implicating IFN-γ/TNF-α in overlapping necroptotic–pyroptotic loss. Importantly, the manner in which cardiomyocytes undergo immune elimination may dictate the downstream inflammatory milieu. Pyroptotic cells form blebbing ‘pyroptotic bodies’ reminiscent of apoptosis, whereas necroptotic cells swell osmotically before catastrophic rupture^73^, releasing distinct sets of damage-associated molecular patterns (e.g., histones)^73^. Cytokine release also differs: IL-33 is liberated chiefly by necroptosis, whereas IL-1β is produced by both necroptosis and pyroptosis^74^. IL-33 supports reparative regulatory T-cell responses that promote healing^75^, whereas IL-1β is linked to higher mortality and recurrent major adverse cardiovascular events^76^. Clarifying how γδ-T cells orchestrate necroptosis, pyroptosis, and potential panoptosis to influence post-MI inflammation will be essential for refining immunomodulatory therapies.

The translational relevance of targeting this stress–ligand axis is further supported by human data: stressed cardiomyocytes express MICA^77^, and elevated levels of soluble MICA (sMICA) following STEMI correlate with larger infarct size, greater cardiomyocyte death, and increased mortality^78^. Notably, MICA has already emerged as a therapeutic target in stress-related cardiometabolic disorders^79–81^, supporting the concept that interruption of IFN-I signalling, and consequently γδ-T cell activation, may represent a novel, lifesaving therapeutic approach for patients with MI. More importantly, our data clearly provide a critical therapeutic window where stressed cardiomyocytes strongly express MCP-1 and RAE-1 and they seem to be innate initiators to recruit γδ-T cells. Along with daily treatment with H-151 for 28 days, only two doses of IFNAR1 mAb given within 48 hours can achieve similar results. The half-life of IgG mAbs is about 21 days^82^ and we have shown that one dose of γδ-T cell-depleting mAb effectively depletes γδ-T cells for 14 days. Therefore, a single dose of IFNAR1 mAb given at the time of myocardial injury should be sufficient to protect stressed cardiomyocytes from RAE-1 NKG2D mediated immune elimination.

Our study has several limitations. First, all experiments were conducted in mice, and the downstream signalling events that follow MI may not be identical in humans. Nonetheless, clinical evidence suggests that the early cardiac stress response is broadly conserved. For example, circulating MCP-1 rises in patients with acute MI^83^, mirroring our murine data. Likewise, the human stress-inducible NKG2D ligand MICA, functionally analogous to murine RAE-1, is upregulated in rejected human hearts^84^□ and has been proposed as an early biomarker of acute MI^65^. MICA is recognised by human γδ-T cells and potently enhances their cytotoxicity^85^, and its induction appears to rely on STING through IFN-α/β- and IRF3-dependent mechanisms^53, 86^. Second, we lacked post-MI cardiac biopsy tissue, preventing direct assessment of cardiomyocyte pyroptosis or necroptosis in patients. Even so, the documented elevations of the nuclear alarmins IL-33 and IL-1β after human MI^87, 88^ are consistent with the activation of necroptotic and pyroptotic death programmes, lending translational support to our murine findings.

In conclusion, our data identify cytotoxic γδ-T cells as key mediators of immunogenic cardiomyocyte death in both PL and IR injury. The trigger is the early release of organelle-derived DNA into the cardiomyocyte cytosol, which activates cGAS–STING signalling. This pathway up-regulates MCP-1, the NKG2D ligand RAE-1, and type□I interferons, while concurrently boosting NKG2D expression on γδ-T cells. The resulting chemotactic and activating signals recruit γδ-T cells to the infarct zone, where cell-to-cell synapses enable delivery of Pfp/GrzB and the synergistic death-promoting cytokines TNF-α and IFN-γ, selectively eliminating stressed cardiomyocytes. Importantly, transient blockade of IFNI signalling with anti-IFNAR antibodies—already approved for lupus therapy—markedly attenuated infarct size in our models, pointing to a readily translatable strategy for improving MI outcomes.

## REFERENCES

1. Bulluck H, Yellon DM, Hausenloy DJ. Reducing myocardial infarct size: challenges and future opportunities. Heart 2016;102(5):341–8.

2. Conrad N, Judge A, Tran J, Mohseni H, Hedgecott D, Crespillo AP, Allison M, Hemingway H, Cleland JG, McMurray JJV, Rahimi K. Temporal trends and patterns in heart failure incidence: a population-based study of 4 million individuals. Lancet 2018;391(10120):572–580.

3. Stewart S, MacIntyre K, Hole DJ, Capewell S, McMurray JJ. More ‘malignant’ than cancer? Five-year survival following a first admission for heart failure. Eur J Heart Fail 2001;3(3):315–22.

4. Steffens S, Van Linthout S, Sluijter JPG, Tocchetti CG, Thum T, Madonna R. Stimulating pro-reparative immune responses to prevent adverse cardiac remodelling: consensus document from the joint 2019 meeting of the ESC Working Groups of cellular biology of the heart and myocardial function. Cardiovasc Res 2020;116(11):1850–1862.

5. Ali M, Pulli B, Courties G, Tricot B, Sebas M, Iwamoto Y, Hilgendorf I, Schob S, Dong A, Zheng W, Skoura A, Kalgukar A, Cortes C, Ruggeri R, Swirski FK, Nahrendorf M, Buckbinder L, Chen JW. Myeloperoxidase Inhibition Improves Ventricular Function and Remodeling After Experimental Myocardial Infarction. JACC Basic Transl Sci 2016;1(7):633–643.

6. Horckmans M, Ring L, Duchene J, Santovito D, Schloss MJ, Drechsler M, Weber C, Soehnlein O, Steffens S. Neutrophils orchestrate post-myocardial infarction healing by polarizing macrophages towards a reparative phenotype. Eur Heart J 2017;38(3):187–197.

7. Yan X, Anzai A, Katsumata Y, Matsuhashi T, Ito K, Endo J, Yamamoto T, Takeshima A, Shinmura K, Shen W, Fukuda K, Sano M. Temporal dynamics of cardiac immune cell accumulation following acute myocardial infarction. J Mol Cell Cardiol 2013;62:24–35.

8. Zouggari Y, Ait-Oufella H, Bonnin P, Simon T, Sage AP, Guerin C, Vilar J, Caligiuri G, Tsiantoulas D, Laurans L, Dumeau E, Kotti S, Bruneval P, Charo IF, Binder CJ, Danchin N, Tedgui A, Tedder TF, Silvestre JS, Mallat Z. B lymphocytes trigger monocyte mobilization and impair heart function after acute myocardial infarction. Nat Med 2013;19(10):1273–80.

9. Hofmann U, Beyersdorf N, Weirather J, Podolskaya A, Bauersachs J, Ertl G, Kerkau T, Frantz S. Activation of CD4+ T lymphocytes improves wound healing and survival after experimental myocardial infarction in mice. Circulation 2012;125(13):1652–63.

10. Weirather J, Hofmann UD, Beyersdorf N, Ramos GC, Vogel B, Frey A, Ertl G, Kerkau T, Frantz S. Foxp3+ CD4+ T cells improve healing after myocardial infarction by modulating monocyte/macrophage differentiation. Circ Res 2014;115(1):55–67.

11. Yan X, Shichita T, Katsumata Y, Matsuhashi T, Ito H, Ito K, Anzai A, Endo J, Tamura Y, Kimura K, Fujita J, Shinmura K, Shen W, Yoshimura A, Fukuda K, Sano M. Deleterious effect of the IL-23/IL-17A axis and gammadeltaT cells on left ventricular remodeling after myocardial infarction. J Am Heart Assoc 2012;1(5):e004408.

12. Knight A, Mackinnon S, Lowdell MW. Human Vdelta1 gamma-delta T cells exert potent specific cytotoxicity against primary multiple myeloma cells. Cytotherapy 2012;14(9):1110–8.

13. von Massow G, Oh S, Lam A, Gustafsson K. Gamma Delta T Cells and Their Involvement in COVID-19 Virus Infections. Front Immunol 2021;12:741218.

14. Kyaw T, Loveland P, Kanellakis P, Cao A, Kallies A, Huang AL, Peter K, Toh BH, Bobik A. Alarmin-activated B cells accelerate murine atherosclerosis after myocardial infarction via plasma cell-immunoglobulin-dependent mechanisms. Eur Heart J 2021;42(9):938–947.

15. Brassington K, Kanellakis P, Cao A, Toh BH, Peter K, Bobik A, Kyaw T. Crosstalk between cytotoxic CD8+ T cells and stressed cardiomyocytes triggers development of interstitial cardiac fibrosis in hypertensive mouse hearts. Front Immunol 2022;13:1040233.

16. Tay C, Liu YH, Kanellakis P, Kallies A, Li Y, Cao A, Hosseini H, Tipping P, Toh BH, Bobik A, Kyaw T. Follicular B Cells Promote Atherosclerosis via T Cell-Mediated Differentiation Into Plasma Cells and Secreting Pathogenic Immunoglobulin G. Arterioscler Thromb Vasc Biol 2018;38(5):e71–e84.

17. Kanellakis P, Pomilio G, Agrotis A, Gao X, Du XJ, Curtis D, Bobik A. Darbepoetin-mediated cardioprotection after myocardial infarction involves multiple mechanisms independent of erythropoietin receptor-common beta-chain heteroreceptor. Br J Pharmacol 2010;160(8):2085–96.

18. Ninh VK, Calcagno DM, Yu JD, Zhang B, Taghdiri N, Sehgal R, Mesfin JM, Chen CJ, Kalhor K, Toomu A, Duran JM, Adler E, Hu J, Zhang K, Christman KL, Fu Z, Bintu B, King KR. Spatially clustered type I interferon responses at injury borderzones. Nature 2024;633(8028):174–181.

19. Kanellakis P, Slater NJ, Du XJ, Bobik A, Curtis DJ. Granulocyte colony-stimulating factor and stem cell factor improve endogenous repair after myocardial infarction. Cardiovasc Res 2006;70(1):117–25.

20. Teichholz LE, Kreulen T, Herman MV, Gorlin R. Problems in echocardiographic volume determinations: echocardiographic-angiographic correlations in the presence of absence of asynergy. Am J Cardiol 1976;37(1):7–11.

21. Fischer MA, Jia L, Edelblum KL. Type I IFN Induces TCR-dependent and - independent Antimicrobial Responses in gammadelta Intraepithelial Lymphocytes. J Immunol 2024;213(9):1380–1391.

22. Takagawa J, Zhang Y, Wong ML, Sievers RE, Kapasi NK, Wang Y, Yeghiazarians Y, Lee RJ, Grossman W, Springer ML. Myocardial infarct size measurement in the mouse chronic infarction model: comparison of area- and length-based approaches. J Appl Physiol (1985) 2007;102(6):2104–11.

23. Cibrian D, Sanchez-Madrid F. CD69: from activation marker to metabolic gatekeeper. Eur J Immunol 2017;47(6):946–953.

24. Ribot JC, deBarros A, Pang DJ, Neves JF, Peperzak V, Roberts SJ, Girardi M, Borst J, Hayday AC, Pennington DJ, Silva-Santos B. CD27 is a thymic determinant of the balance between interferon-gamma- and interleukin 17-producing gammadelta T cell subsets. Nat Immunol 2009;10(4):427–36.

25. Karki R, Sharma BR, Tuladhar S, Williams EP, Zalduondo L, Samir P, Zheng M, Sundaram B, Banoth B, Malireddi RKS, Schreiner P, Neale G, Vogel P, Webby R, Jonsson CB, Kanneganti TD. Synergism of TNF-alpha and IFN-gamma Triggers Inflammatory Cell Death, Tissue Damage, and Mortality in SARS-CoV-2 Infection and Cytokine Shock Syndromes Cell 2021;184(1):149–168 e17.

26. van Loo G, Bertrand MJM. Death by TNF: a road to inflammation. Nat Rev Immunol 2023;23(5):289–303.

27. Zhou Z, He H, Wang K, Shi X, Wang Y, Su Y, Wang Y, Li D, Liu W, Zhang Y, Shen L, Han W, Shen L, Ding J, Shao F. Granzyme A from cytotoxic lymphocytes cleaves GSDMB to trigger pyroptosis in target cells. Science 2020;368(>6494).

28. Zou Y, Li L, Li Y, Chen S, Xie X, Jin X, Wang X, Ma C, Fan G, Wang W. Restoring Cardiac Functions after Myocardial Infarction-Ischemia/Reperfusion via an Exosome Anchoring Conductive Hydrogel. ACS Appl Mater Interfaces 2021;13(48):56892–56908.

29. Chen L, Woo M, Hakem R, Miller RG. Perforin-dependent activation-induced cell death acts through caspase 3 but not through caspases 8 or 9. Eur J Immunol 2003;33(3):769–78.

30. Prabhu SD, Frangogiannis NG. The Biological Basis for Cardiac Repair After Myocardial Infarction: From Inflammation to Fibrosis. Circ Res 2016;119(1):91–112.

31. Gong W, Fang P, Leng M, Shi Y. Promoting GSDME expression through DNA demethylation to increase chemosensitivity of breast cancer MCF-7 / Taxol cells. PLoS One 2023;18(3):e0282244.

32. Liu OH, Kiema M, Beter M, Yla-Herttuala S, Laakkonen JP, Kaikkonen MU. Hypoxia-Mediated Regulation of Histone Demethylases Affects Angiogenesis-Associated Functions in Endothelial Cells. Arterioscler Thromb Vasc Biol 2020;40(11):2665–2677.

33. Ferreira FM, Palle P, Vom Berg J, Prajwal P, Laman JD, Buch T. Bone marrow chimeras-a vital tool in basic and translational research. J Mol Med (Berl) 2019;97(7):889–896.

34. Chang HF, Schirra C, Ninov M, Hahn U, Ravichandran K, Krause E, Becherer U, Balint S, Harkiolaki M, Urlaub H, Valitutti S, Baldari CT, Dustin ML, Jahn R, Rettig J. Identification of distinct cytotoxic granules as the origin of supramolecular attack particles in T lymphocytes. Nat Commun 2022;13(1):1029.

35. Knuth AK, Rosler S, Schenk B, Kowald L, van Wijk SJL, Fulda S. Interferons Transcriptionally Up-Regulate MLKL Expression in Cancer Cells. Neoplasia 2019;21(1):74–81.

36. Metkar SS, Wang B, Aguilar-Santelises M, Raja SM, Uhlin-Hansen L, Podack E, Trapani JA, Froelich CJ. Cytotoxic cell granule-mediated apoptosis: perforin delivers granzyme B-serglycin complexes into target cells without plasma membrane pore formation. Immunity 2002;16(3):417–28.

37. Xu C, Wu J, Wu Y, Ren Z, Yao Y, Chen G, Fang EF, Noh JH, Liu YU, Wei L, Chen X, Sima J. TNF-alpha-dependent neuronal necroptosis regulated in Alzheimer’s disease by coordination of RIPK1-p62 complex with autophagic UVRAG. Theranostics 2021;11(19):9452–9469.

38. Wang H, Sun L, Su L, Rizo J, Liu L, Wang LF, Wang FS, Wang X. Mixed lineage kinase domain-like protein MLKL causes necrotic membrane disruption upon phosphorylation by RIP3. Mol Cell 2014;54(1):133–146.

39. Gautam A, Boyd DF, Nikhar S, Zhang T, Siokas I, Van de Velde LA, Gaevert J, Meliopoulos V, Thapa B, Rodriguez DA, Cai KQ, Yin C, Schnepf D, Beer J, DeAntoneo C, Williams RM, Shubina M, Livingston B, Zhang D, Andrake MD, Lee S, Boda R, Duddupudi AL, Crawford JC, Vogel P, Loch C, Schwemmle M, Fritz LC, Schultz-Cherry S, Green DR, Cuny GD, Thomas PG, Degterev A, Balachandran S. Necroptosis blockade prevents lung injury in severe influenza. Nature 2024;628(8009):835–843.

40. Robinson N, Ganesan R, Hegedus C, Kovacs K, Kufer TA, Virag L. Programmed necrotic cell death of macrophages: Focus on pyroptosis, necroptosis, and parthanatos. Redox Biol 2019;26:101239.

41. Nguyen LN, Kanneganti TD. PANoptosis in Viral Infection: The Missing Puzzle Piece in the Cell Death Field. J Mol Biol 2022;434(4):167249.

42. Lee S, Karki R, Wang Y, Nguyen LN, Kalathur RC, Kanneganti TD. AIM2 forms a complex with pyrin and ZBP1 to drive PANoptosis and host defence. Nature 2021;597(7876):415–419.

43. Paludan SR, Reinert LS, Hornung V. DNA-stimulated cell death: implications for host defence, inflammatory diseases and cancer. Nat Rev Immunol 2019;19(3):141–153.

44. Kabelitz D, Serrano R, Kouakanou L, Peters C, Kalyan S. Cancer immunotherapy with gammadelta T cells: many paths ahead of us. Cell Mol Immunol 2020;17(9):925–939.

45. Miller KN, Victorelli SG, Salmonowicz H, Dasgupta N, Liu T, Passos JF, Adams PD. Cytoplasmic DNA: sources, sensing, and role in aging and disease. Cell 2021;184(22):5506–5526.

46. Napolitano G, Fasciolo G, Venditti P. Mitochondrial Management of Reactive Oxygen Species. Antioxidants (Basel) 2021;10(11).

47. Tian Y, Charles EJ, Yan Z, Wu D, French BA, Kron IL, Yang Z. The myocardial infarct-exacerbating effect of cell-free DNA is mediated by the high-mobility group box 1-receptor for advanced glycation end products-Toll-like receptor 9 pathway. J Thorac Cardiovasc Surg 2019;157(6):2256–2269 e3.

48. Dong J, Holthaus D, Peters C, Koster S, Ehsani M, Quevedo-Olmos A, Berger H, Zarobkiewicz M, Mangler M, Gurumurthy RK, Hedemann N, Chumduri C, Kabelitz D, Meyer TF. gammadelta T cell-mediated cytotoxicity against patient-derived healthy and cancer cervical organoids. Front Immunol 2023;14:1281646.

49. Li W, Li Y, Zhao J, Liao J, Wen W, Chen Y, Cui H. Release of damaged mitochondrial DNA: A novel factor in stimulating inflammatory response. Pathol Res Pract 2024;258:155330.

50. Abdullah A, Zhang M, Frugier T, Bedoui S, Taylor JM, Crack PJ. STING-mediated type-I interferons contribute to the neuroinflammatory process and detrimental effects following traumatic brain injury. J Neuroinflammation 2018;15(1):323.

51. Lanng KRB, Lauridsen EL, Jakobsen MR. The balance of STING signaling orchestrates immunity in cancer. Nat Immunol 2024;25(7):1144–1157.

52. Rothenfusser S, Hornung V, Krug A, Towarowski A, Krieg AM, Endres S, Hartmann G. Distinct CpG oligonucleotide sequences activate human gamma delta T cells via interferon-alpha/-beta. Eur J Immunol 2001;31(12):3525–34.

53. Zhang C, Zhang J, Sun R, Feng J, Wei H, Tian Z. Opposing effect of IFNgamma and IFNalpha on expression of NKG2 receptors: negative regulation of IFNgamma on NK cells. Int Immunopharmacol 2005;5(6):1057–67.

54. Agerholm R, Kadekar D, Rizk J, Bekiaris V. Type I interferon supports gammadelta T-cell homeostasis and immunity through direct and indirect receptor signaling in mice. Eur J Immunol 2021;51(12):3186–3193.

55. Rieckmann M, Delgobo M, Gaal C, Buchner L, Steinau P, Reshef D, Gil-Cruz C, Horst ENT, Kircher M, Reiter T, Heinze KG, Niessen HW, Krijnen PA, van der Laan AM, Piek JJ, Koch C, Wester HJ, Lapa C, Bauer WR, Ludewig B, Friedman N, Frantz S, Hofmann U, Ramos GC. Myocardial infarction triggers cardioprotective antigen-specific T helper cell responses. J Clin Invest 2019;129(11):4922–4936.

56. Iannello A, Raulet DH. Immune surveillance of unhealthy cells by natural killer cells. Cold Spring Harb Symp Quant Biol 2013;78:249–57.

57. Ribot JC, Lopes N, Silva-Santos B. gammadelta T cells in tissue physiology and surveillance. Nat Rev Immunol 2021;21(4):221–232.

58. Lodrini AM, Goumans MJ. Cardiomyocytes Cellular Phenotypes After Myocardial Infarction. Front Cardiovasc Med 2021;8:750510.

59. Riley JS, Tait SW. Mitochondrial DNA in inflammation and immunity. EMBO Rep 2020;21(4):e49799.

60. Domizio JD, Gulen MF, Saidoune F, Thacker VV, Yatim A, Sharma K, Nass T, Guenova E, Schaller M, Conrad C, Goepfert C, de Leval L, Garnier CV, Berezowska S, Dubois A, Gilliet M, Ablasser A. The cGAS-STING pathway drives type I IFN immunopathology in COVID-19. Nature 2022;603(7899):145–151.

61. Boguszewska K, Szewczuk M, Kazmierczak-Baranska J, Karwowski BT. The Similarities between Human Mitochondria and Bacteria in the Context of Structure, Genome, and Base Excision Repair System. Molecules 2020;25(12).

62. Cheung GT, Siow YL, O K. Homocysteine stimulates monocyte chemoattractant protein-1 expression in mesangial cells via NF-kappaB activation. Can J Physiol Pharmacol 2008;86(3):88–96.

63. Hopfner KP, Hornung V. Molecular mechanisms and cellular functions of cGAS-STING signalling. Nat Rev Mol Cell Biol 2020;21(9):501–521.

64. Solis M, Goubau D, Romieu-Mourez R, Genin P, Civas A, Hiscott J. Distinct functions of IRF-3 and IRF-7 in IFN-alpha gene regulation and control of anti-tumor activity in primary macrophages. Biochem Pharmacol 2006;72(11):1469–76.

65. Fu C, Shi Y, Yao Z. sMICA as novel and early predictors for acute myocardial infarction. Eur J Med Res 2016;21(1):25.

66. Lam AR, Bert NL, Ho SS, Shen YJ, Tang LF, Xiong GM, Croxford JL, Koo CX, Ishii KJ, Akira S, Raulet DH, Gasser S. RAE1 ligands for the NKG2D receptor are regulated by STING-dependent DNA sensor pathways in lymphoma. Cancer Res 2014;74(8):2193–2203.

67. Alkhayer R, Ponath V, Frech M, Adhikary T, Graumann J, Neubauer A, von Strandmann EP. KLF4-mediated upregulation of the NKG2D ligand MICA in acute myeloid leukemia: a novel therapeutic target identified by enChIP. Cell Commun Signal 2023;21(1):94.

68. Hosomi S, Grootjans J, Tschurtschenthaler M, Krupka N, Matute JD, Flak MB, Martinez-Naves E, Gomez Del Moral M, Glickman JN, Ohira M, Lanier LL, Kaser A, Blumberg R. Intestinal epithelial cell endoplasmic reticulum stress promotes MULT1 up-regulation and NKG2D-mediated inflammation J Exp Med 2017;214(10):2985–2997.

69. Jones AB, Rocco A, Lamb LS, Friedman GK, Hjelmeland AB. Regulation of NKG2D Stress Ligands and Its Relevance in Cancer Progression. Cancers (Basel) 2022;14(9).

70. Jiang M, Qi L, Li L, Li Y. The caspase-3/GSDME signal pathway as a switch between apoptosis and pyroptosis in cancer. Cell Death Discov 2020;6:112.

71. De Schutter E, Croes L, Ibrahim J, Pauwels P, Op de Beeck K, Vandenabeele P, Van Camp G. GSDME and its role in cancer: From behind the scenes to the front of the stage. Int J Cancer 2021;148(12):2872–2883.

72. Shi H, Gao Y, Dong Z, Yang J, Gao R, Li X, Zhang S, Ma L, Sun X, Wang Z, Zhang F, Hu K, Sun A, Ge J. GSDMD-Mediated Cardiomyocyte Pyroptosis Promotes Myocardial I/R Injury. Circ Res 2021;129(3):383–396.

73. Chen Z, Chen C, Lai K, Wu C, Wu F, Chen Z, Ye K, Xie J, Ma H, Chen H, Wang Y, Xu Y. GSDMD and GSDME synergy in the transition of acute kidney injury to chronic kidney disease. Nephrol Dial Transplant 2024;39(8):1344–1359.

74. Chen X, He WT, Hu L, Li J, Fang Y, Wang X, Xu X, Wang Z, Huang K, Han J. Pyroptosis is driven by non-selective gasdermin-D pore and its morphology is different from MLKL channel-mediated necroptosis. Cell Res 2016;26(9):1007–20.

75. Frank D, Vince JE. Pyroptosis versus necroptosis: similarities, differences, and crosstalk. Cell Death Differ 2019;26(1):99–114.

76. Xia N, Lu Y, Gu M, Li N, Liu M, Jiao J, Zhu Z, Li J, Li D, Tang T, Lv B, Nie S, Zhang M, Liao M, Liao Y, Yang X, Cheng X. A Unique Population of Regulatory T Cells in Heart Potentiates Cardiac Protection From Myocardial Infarction. Circulation 2020;142(20):1956–1973.

77. Wei L, Lu J, Feng L, Long D, Shan J, Li S, Li Y. HIF-1alpha accumulation upregulates MICA and MICB expression on human cardiomyocytes and enhances NK cell cytotoxicity during hypoxia-reoxygenation. Life Sci 2010;87(3-4):111–9.

78. Haohan S, Pussadhamma B, Jumnainsong A, Leuangwatthananon W, Makarawate P, Leelayuwat C, Komanasin N. Association of Major Histocompatibility Complex Class I Related Chain A/B Positive Microparticles with Acute Myocardial Infarction and Disease Severity. Diagnostics (Basel) 2020;10(10).

79. Van Belle TL, Ling E, Haase C, Bresson D, Urso B, von Herrath MG. NKG2D blockade facilitates diabetes prevention by antigen-specific Tregs in a virus-induced model of diabetes. J Autoimmun 2013;40:66–73.

80. Xia M, Guerra N, Sukhova GK, Yang K, Miller CK, Shi GP, Raulet DH, Xiong N. Immune activation resulting from NKG2D/ligand interaction promotes atherosclerosis. Circulation 2011;124(25):2933–43.

81. Gonzalez S, Lopez-Soto A, Suarez-Alvarez B, Lopez-Vazquez A, Lopez-Larrea C. NKG2D ligands: key targets of the immune response. Trends Immunol 2008;29(8):397–403.

82. Baldwin WM, 3rd, Valujskikh A, Fairchild RL. The neonatal Fc receptor: Key to homeostasic control of IgG and IgG-related biopharmaceuticals. Am J Transplant 2019;19(7):1881–1887.

83. Matsumori A, Furukawa Y, Hashimoto T, Yoshida A, Ono K, Shioi T, Okada M, Iwasaki A, Nishio R, Matsushima K, Sasayama S. Plasma levels of the monocyte chemotactic and activating factor/monocyte chemoattractant protein-1 are elevated in patients with acute myocardial infarction. J Mol Cell Cardiol 1997;29(1):419–23.

84. Suarez-Alvarez B, Lopez-Vazquez A, Gonzalez MZ, Fdez-Morera JL, Diaz-Molina B, Blanco-Gelaz MA, Pascual D, Martinez-Borra J, Muro M, Alvarez-Lopez MR, Lopez-Larrea C. The relationship of anti-MICA antibodies and MICA expression with heart allograft rejection. Am J Transplant 2007;7(7):1842–8.

85. Li J, Cui L, He W. Distinct pattern of human Vdelta1 gammadelta T cells recognizing MICA. Cell Mol Immunol 2005;2(4):253–8.

86. Raulet DH, Gasser S, Gowen BG, Deng W, Jung H. Regulation of ligands for the NKG2D activating receptor. Annu Rev Immunol 2013;31:413–41.

87. Silvain J, Kerneis M, Zeitouni M, Lattuca B, Galier S, Brugier D, Mertens E, Procopi N, Suc G, Salloum T, Frisdal E, Le Goff W, Collet JP, Vicaut E, Lesnik P, Montalescot G, Guerin M. Interleukin-1beta and Risk of Premature Death in Patients With Myocardial Infarction. J Am Coll Cardiol 2020;76(15):1763–1773.

88. Xing J, Liu J, Geng T. Predictive values of sST2 and IL-33 for heart failure in patients with acute myocardial infarction. Exp Biol Med (Maywood) 2021;246(23):2480–2486.

